# LyTS: A Lysosome Localized Complex of TMEM192 and STK11IP

**DOI:** 10.1101/2021.08.27.457973

**Authors:** B. Angarola, F. Frölich, S.M. Ferguson

## Abstract

The degradative and signaling functions of lysosomes are dependent on numerous peripherally associated proteins. Targeting of lysosomes to sites of need is controlled by adaptors that link lysosomes to both dynein and kinesin motors. SKIP is one such adaptor that promotes microtubule plus-end-directed movement through its interaction with Arl8 on the lysosome surface and kinesin-1. Sequence homology between SKIP and STK11IP (also known as LIP1) led us to investigate a potential role for STK11IP at lysosomes. After first establishing that STK11IP localizes to lysosomes, we identified TMEM192, an abundant lysosomal integral membrane protein, as the major binding partner of STK11IP and demonstrated that STK11IP depends on TMEM192 for both its lysosome localization as well as its stability. Depletion studies furthermore support a role for these proteins in the control of lysosome homeostasis. Collectively, these new results define a lysosome localized complex of TMEM192 and STK11IP that we have named LyTS (“lights”).

## Introduction

As recipients of material delivered both by the endocytic and autophagy pathways, lysosomes support multiple critical cellular processes including: clearance of defective organelles and proteins, degrading harmful pathogens, modulating signaling through the turnover of activated receptors, and digesting endocytosed nutrients (Ballabio and Bonifacino, 2019; Lim and Zoncu, 2016; Liu and Sabatini, 2020; Luzio et al., 2007). In addition to the degradative and nutrient recycling functions carried out by luminal hydrolases, integral membrane transporters, and ion channels, lysosomes depend on a large collection of proteins that are recruited to their cytoplasmic surface to support signaling as well as to control their positioning within cells (Ferguson, 2015; Liu and Sabatini, 2020; Saftig and Puertollano, 2021). The movement of lysosomes depends on adaptors that link lysosomes to either dynein or kinesin motors (Bonifacino and Neefjes, 2017; Ferguson, 2018). One important mechanism for linking lysosomes to kinesin is via an adaptor known as SKIP (SifA and Kinesin Interacting Protein) that forms a bridge between Arl8, a small GTPase enriched on lysosomes, and kinesin 1 and thus promotes the movement of lysosomes towards the plus ends of microtubules at the cell periphery (Farias et al., 2017; Hofmann and Munro, 2006; Keren-Kaplan and Bonifacino, 2021; Korolchuk et al., 2011; Pu et al., 2015; Rosa-Ferreira and Munro, 2011; Rosa-Ferreira et al., 2018)..

Given the importance of lysosome subcellular positioning and the established role for SKIP in this process, we investigated serine/threonine kinase 11-interactingprotein (STK11IP; also known as LIP1), a protein that was proposed to share homology with SKIP, but whose subcellular functions and site of action was not well established. Our results show that STK11IP is robustly recruited to lysosomes through an interaction with transmembrane protein 192 (TMEM192) and depends on this interaction for its stability. We further demonstrate that depletion of either TMEM192 or STK11IP results in similar defects in mTORC1 signaling, a process that is broadly sensitive to perturbations to lysosome homeostasis.

## Results and Discussion

STK11IP was originally identified as a novel interacting protein of serine/threonine kinase 11/Liver Kinase B1 (STK11/LKB1) in a yeast two hybrid screen (Smith et al., 2001). In addition to the proposed interaction with STK11, STK11IP was also reported to interact with SMAD4 and Toll Like Receptors 4 and 9 (TLR4 and TLR9) (Moren et al., 2011; Smith et al., 2001). However, despite these efforts to characterize the STK11IP protein, its subcellular site of action and physiological functions remained largely unclear. STK11IP is a 1099 amino acid protein with an amino-terminal domain containing of leucine-rich repeats (LRR) and a carboxyl-terminal domain (CTD) of unknown function (Fig. 1A). One potential clue regarding STK11IP function is that the CTD of LIP was previously reported to share limited homology with two proteins: Nischarin and SKIP (SifA and kinesin-interacting protein) (Rosa-Ferreira and Munro, 2011). This C-terminal region of SKIP is now known to contain 3 PH domains (Keren-Kaplan and Bonifacino, 2021). Recent structural predictions from Alphafold2 also define putative PH domains in the C-terminus of STK11IP (Tunyasuvunakool et al., 2021). Nischarin was also implicated in endosome maturation (Kuijl et al., 2013). Based on predicted structural similarities between SKIP, Nischarin, and STK11IP and the known roles for SKIP and Nischarin in the endolysosomal pathway, we thus investigated STK11IP subcellular localization.

**Figure 1.**
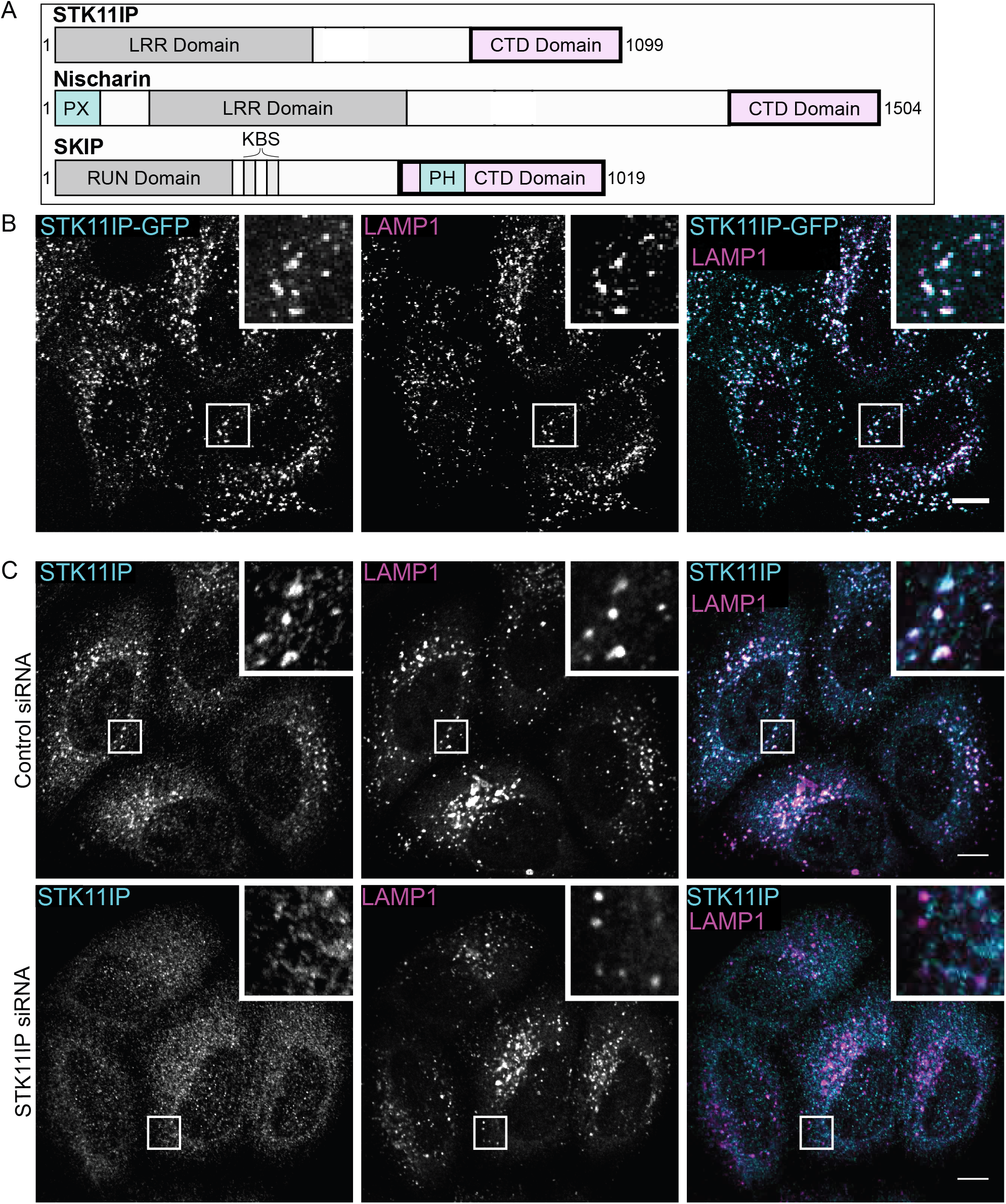
STK11IP localizes to lysosomes. **(A)** Simple schematic domain organization of STK11IP, Nischarin and SKIP proteins. Highlighted regions include: LRR (Leucine Rich Repeat), CTD Domain (Carboxyl-Terminal Domain), PX (Phox homology domain), RUN domain, kinesin binding sites (KBS) and PH (Pleckstrin Homology) domain. **(B)** Representative images of HeLa cells transfected with STK11IP-GFP with anti-GFP and anti-LAMP1 immunofluorescence. **(C)** Representative immunofluorescence images of endogenous STK11IP localization using anti-STK11IP anti-LAMP1 antibodies in HeLa cells treated with Control or STK11IP siRNA. Scale bars, 10 μm.

### STK11IP localizes to lysosomes

Although SKIP is localized to lysosomes via an interaction with Arl8 that is mediated by its RUN domain and Nischarin binds endosomes via an interaction between its PX domain and PI3P, STK11IP lacks either of these putative endolysomal targeting domains (Kuijl et al., 2013; Rosa-Ferreira and Munro, 2011). Nonetheless, analysis of a previously published proteomic analysis of purified lysosomes, suggested that STK11IP might be present at lysosomes (Schroder et al., 2007). However, another study had reported that STK11IP was cytoplasmic (Smith et al., 2001). To directly investigate STK11IP subcellular localization, we expressed STK11IP-GFP in Hela cells and observed that it exhibited a highly punctate localization pattern that co-localized extensively with the late endosome and lysosome marker, lysosome associated membrane protein 1 (LAMP1; Fig. 1B). We also performed immunofluorescence with an antibody that recognizes the endogenous STK11IP1 and once again found that was enriched on LAMP1-positive compartments (Figure 1C). The specificity of this LAMP1-colocalized immunofluorescence signal for STK11IP was demonstrated by the fact that it was absent following siRNA-mediated depletion of STK11IP (Figure 1D).

### An interaction with TMEM192 is important for STK11IP localization and stability

To identify novel proteins that mediate the lysosomal localization of STK11IP, we performed anti-GFP immunoprecipitations from HeLa cells that stably expressed STK11IP-GFP and analyzed these samples with label free quantitative mass spectrometry (LC-MS/MS). As expected, STK11IP was the most abundant and enriched protein in the sample (Figure 2A). Interestingly, TMEM192, an integral membrane protein of lysosomes (Schroder et al., 2010), was identified as a prominent STK11IP interacting partner (Figure 2A). We validated this interaction with co-immunoprecipitation and immunoblotting experiments (Figure 2B). The importance of the STK11IP-TMEM192 interaction was further established by observations that siRNA mediated depletion of TMEM192 led to a major loss of STK11IP which indicates that the STK11IP is unstable in the absence of its interaction with TMEM192 (Figure 2C). We furthermore observed that STK11IP and TMEM192 colocalize on lysosomes via immunofluorescence microscopy (Figure 2D) and that the STK11IP protein that remained following TMEM192 depletion no longer localized to lysosomes (Figure 2E). These results strongly suggest that STK11IP and TMEM192 are in a stable complex at the surface of lysosomes. We did not detect enrichment for STK11 in our STK11IP immunoprecipitations. However, as HeLa cells may lack functional STK11, our model system may not have been suitable for capturing this previously reported interaction (McCabe et al., 2010). Thus, while we cannot rule out a role for STK11-STK11IP interactions in other contexts, we conclude that the interaction between STK11IP and TMEM192 is critical for STK11IP subcellular localization and stability and is independent of the previously reported STK11 interaction.

**Figure 2.**
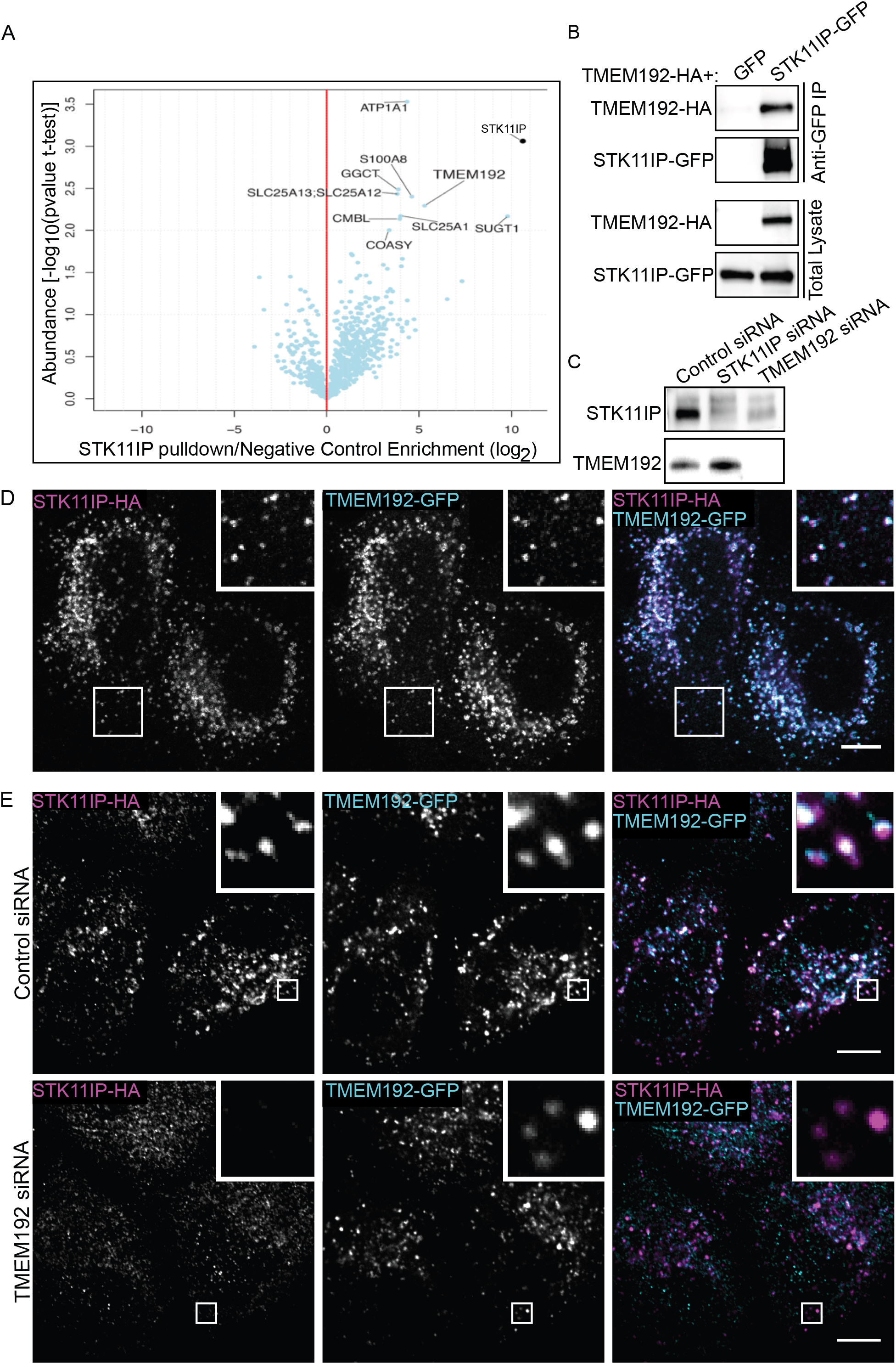
STK11IP is recruited to lysosomes via an interaction with TMEM192. **(A)** Volcano plot summarizing the STK11IP-GFP interacting proteins identified by label-free mass spectrometry analysis of anti-GFP immunoprecipitates. **(B)** Western blot analysis of anti-GFP pulldowns of HeLa cells transfected with TMEM192-HA + GFP or STK11IP-GFP. **(C)** Western blot analysis of STK11IP and TMEM192 levels in HeLa cells treated with control, STK11IP, and TMEM192 siRNAs. **(D)** Representative images of HeLa cells transfected with STK11IP-HA and TMEM192-GFP with detection by anti-HA and anti-GFP immunofluorescence. **(E)** Representative immunofluorescence images of endogenous STK11IP localization to lysosomes using anti-STK11IP anti-LAMP1 antibodies in HeLa cells treated with control or TMEM192 siRNA. Scale bars, 10 μm.

### Mapping interactions between TMEM192 and STK11IP

TMEM192 is a 271 amino acid lysosome membrane protein with 4 transmembrane domains that contains cytosolic N and C termini (Figure 3A)(Behnke et al., 2011; Schroder et al., 2010). To identify the regions within STK11IP and TMEM192 that mediate their interaction, we designed a series of truncation mutants. When mutating TMEM192, we focused on the cytosolic N and C termini of TMEM192, as these were the largest candidate regions within TMEM192 for interactions with STK11IP at the cytosolic face of the lysosome. GFP-TMEM192^ΔN40^ (lacking the N terminus) weakly interacted with STK11IP (Figure 3B). However, the interpretation of this result was confounded by the fact that this deletion removes the dileucine motifs that are critical for its lysosomal localization (Behnke et al., 2011; Braulke and Bonifacino, 2009). In contrast, deletions within the C-terminus of TMEM192 (TMEM192^ΔC34^-GFP and TMEM192^ΔC73-^GFP) abolished the interaction (Figure 3B). As the localization of the TMEM192^ΔC34^-GFP and TMEM192^ΔC73^-GFP deletion mutants was still lysosomal (not shown), we conclude that the the C-terminus of TMEM192 likely mediates the interaction with STK11IP (Figure 3B). Recent TMEM192 structural predictions from AlphaFold2 (Tunyasuvunakool et al., 2021) propose that the most prominent feature within the TMEM192 C-terminus is an alpha helix comprised by amino acids 228-265. More focused mutagenesis based on this new information may help to resolve the precise determinants within TMEM192 that supports its ability to interact with STK11IP.

**Figure 3.**
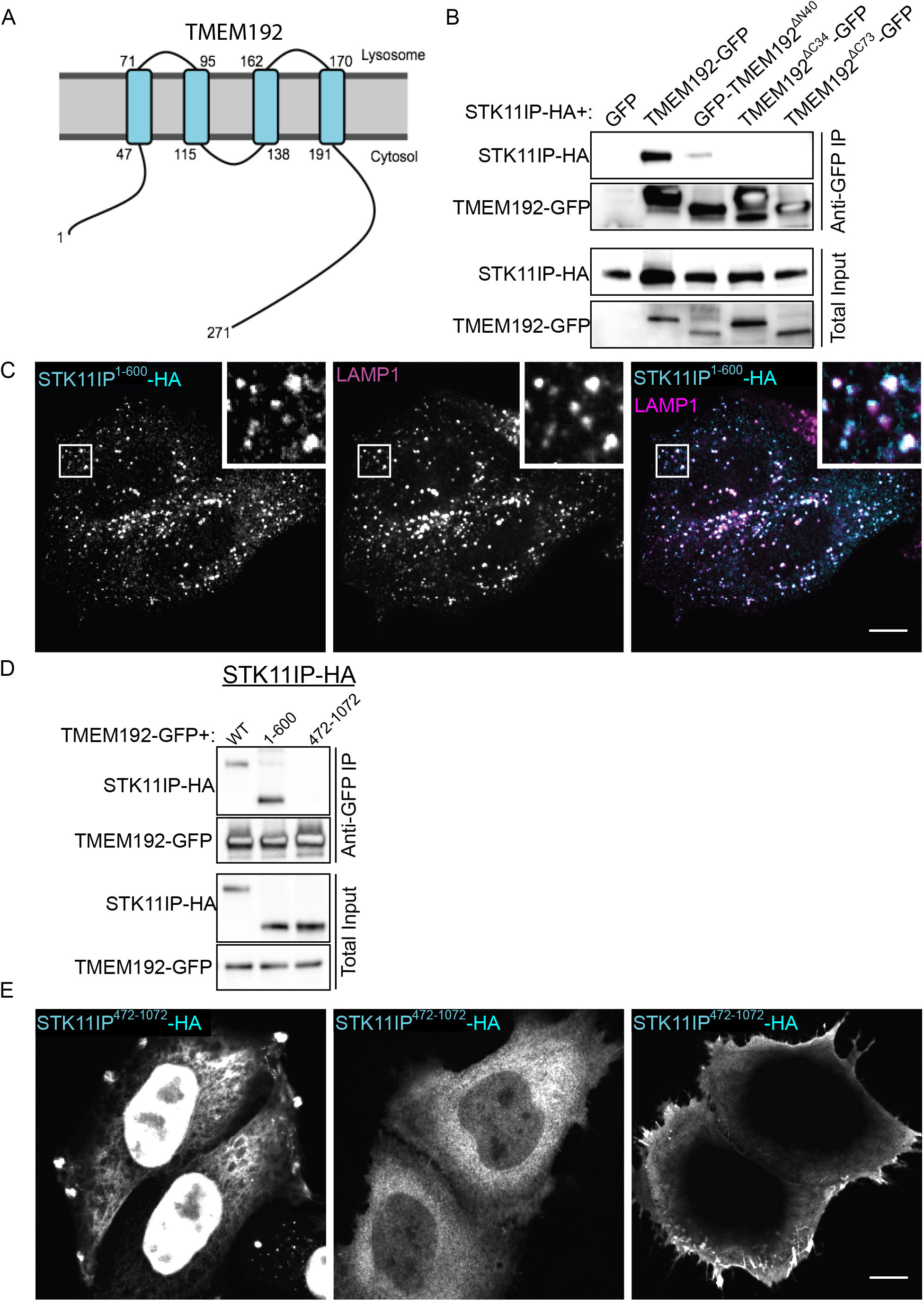
The C-terminus of TMEM192 interacts with the first 600 amino acids of STK11IP. **(A)** Schematic diagram of the topology of TMEM192. TMEM192 has 4 transmembrane domains and both the N-an C-termini are cytoplasmic. **(B)** Western blot analysis of anti-GFP pulldowns of HeLa cells transfected with STK11IP-HA + GFP, TMEM192-GPF, GFP-TMEM192^ΔN40^, TMEM192^ΔC34^-GFP, or TMEM192^ΔC73^-GFP. **(C)** Anti-HA and anti-LAMP1 immunofluorescence in representative images of HeLa cells expressing STK11IP^1-600^-HA. **(D)** Immunoblot analysis of anti-GFP pulldowns of HeLa cells expressing TMEM192-GFP + STK11IP-HA, STK11IP^1-600^-HA, or HA-STK11IP^472-1072^. **(E)** Examples of anti-HA immunofluorescence in HeLa cells expressing HA-STK11IP^472-1072^ that summarize the variety of distinct localization patterns that were observed for this protein. Scale bars, 10 μm.

To better understand which parts of STK11IP are critical for its lysosomal localization, we created two truncated versions of STK11IP: STK11IP^1-600^-HA and STK11IP^472-1072^-HA. These mutations were made based on predictions that the region between amino acids 472 and 600 might contain a coiled-coil that could support STK11IP dimerization/oligomerization (Lupas et al., 1991; Moren et al., 2011). Interestingly, STK11IP^1-600^-HA localized lysosomes (Figure 3C). We furthermore found that TMEM192 still interacts with the STK11IP^1-600^ fragment (Figure 3D).

Conversely, the HA-STK11IP^472-1072^ mutant did not interact with TMEM192 and was not lysosome-localized (Figure 3D and E). Interestingly, this protein had multiple distinct patterns of subcellular localization that varied from cell-to-cell (Figure 3E). This variable localization might reflect multiple C-terminal interacting proteins or a complex ability to bind various intracellular membranes. Using the HHPred protein structure prediction server, we identified a predicted Pleckstrin Homology (PH) domain in the CTD of STK11IP (Fidler et al., 2016). The presence of PH domains within the STK11IP C-terminus were also very recently predicted by Alphafold2 (Tunyasuvunakool et al., 2021). PH domains can bind phosphoinositides and therefore can aid in the recruitment of proteins to membranes (Lemmon, 2007). Therefore, it is possible that STK11IP binds to the lysosome by a bipartite mechanism through the interaction with TMEM192 and via a PH domain with modest membrane affinity/selectivity. However, additional roles for PH domains within STK11IP are possible. For example, C-terminal PH domains within SKIP were recently proposed to mediate intramolecular interactions that inhibit SKIP function in the absence of Arl8 binding (Keren-Kaplan and Bonifacino, 2021).

### Potential function of a STK11IP-TMEM192 complex

TMEM192 was reported to regulate autophagy and apoptosis in hepatoma cells and cervical cells (Liu et al., 2012; Shyu et al., 2016). Consistent with these observations and a well-established role for mTORC1 signaling as a negative regulator of autophagy (Liu and Sabatini, 2020), we found that knocking down either STK11IP or TMEM192 resulted in a reduction in the phosphorylation of ribosomal protein S6, a major downstream target of the mTORC1 pathway (Figure 4A and 4B). Although these results support a functional impact of TMEM192-STK11IP depletion on mTORC1 signaling from lysosomes, the directness of the link to the mTORC1 machinery remains to be elucidated.

**Figure 4.**
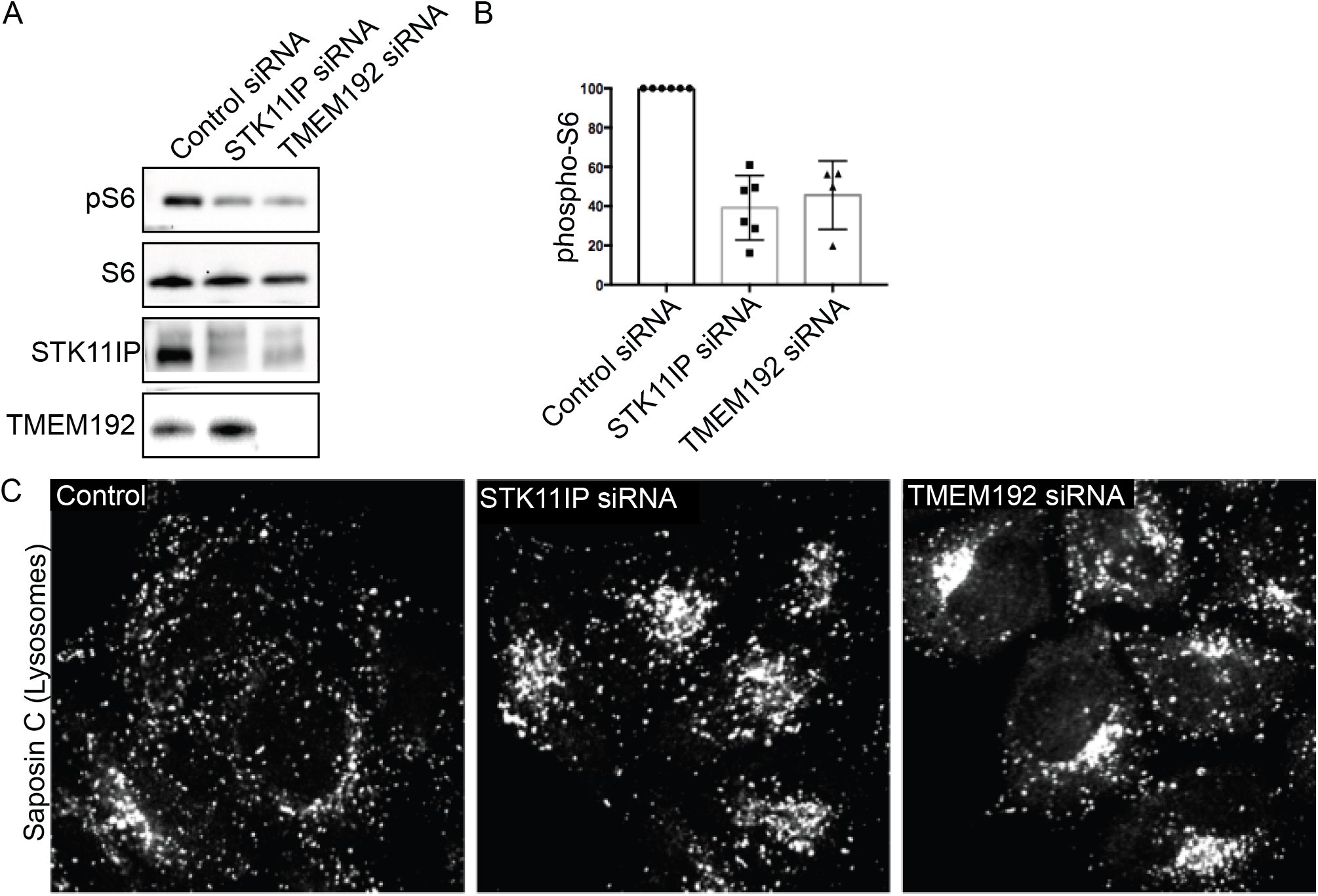
Potential functional roles for TMEM192-STK11IP complex at lysosomes. **(A)** Immunoblot analysis of ribososmal protein S6 phosphorylation in HelaM cells treated with the indicated siRNAs. STK11IP and TMEM192 blots are duplicated from Figure 2C as the same samples were analyzed in each of these experiments. **(B)** Quantification of S6 phosphorylation under the indicated conditions. Phospho-S6 levels were normalized to total S6 levels in each sample. **(C)** Representative immunofluorescence images of saposin C (lysosome marker) localization in cells treated with the indicated siRNAs.

Given the role for SKIP in promoting kinesin-dependent movement of lysosomes, we also examined the impact of STKI11IP and TMEM192 depletion on lysosome subcellular positioning and found a modest increase in perinuclear accumulation lysosomes (labeled with Saposin C antibodies; Figure 4C). However, we also observed that overexpressing STK11IP and TMEM192 did not reposition lysosomes towards the cell periphery (Figure 2D). In contrast, overexpressing Arl8 and/or SKIP dramatically redistributes lysosomes (Keren-Kaplan and Bonifacino, 2021; Rosa-Ferreira and Munro, 2011).

Interestingly, TMEM192 KO mice were reported to have no detectable lysosome defects (Nguyen et al., 2017). This finding is at odds with the robust localization of TMEM192 to lysosomes and could reflect either the need for more specific assays; compensatory mechanisms; and/or redundant pathways that are engaged following long-term loss of TMEM192. Interestingly, it was very recently reported that STK11IP is an mTORC1 substrate and autophagy inhibitor (Zi et al., 2021). Given the major role that we have uncovered for the interaction between TMEM192 and STK11IP in controlling both the abundance and subcellular localization of STK11IP, it will be of interest to integrate the LyTS complex into future studies that define physiological functions of both the STK11IP and TMEM192 proteins.

As an increasing number of investigators are using over-expressed epitope tagged TMEM192 to purify lysosomes (Abu-Remaileh et al., 2017; Singh et al., 2020), it will be important for users of this tool to take into account how over-expression of TMEM192 and its interaction with STK11IP might impact results from experiments that use this strategy for lysosome enrichment.

In conclusion, we have identified a novel lysosome-localized complex of TMEM192-STK11IP that we call LyTS (“lights”). These efforts form a foundation for future studies into the physiological functions of these proteins.

## Materials and Methods

### Cell Culture

HeLa M cells (kindly provided by P. De Camilli, Yale University, New Haven) were grown in high-glucose Dulbecco’s Modified Eagle’s medium (DMEM) with L-glutamine, 10% fetal bovine serum, and 1% penicillin/streptomycin supplement (ThermoFisher Scientific, Waltham, MA and Corning, Corning, NY). For plasmid transfections, 500 ng of plasmid DNA, 1.5 μl of Fugene 6 transfection reagent (Promega, Madison, WI) and 100 μl of OptiMEM (ThermoFisher Scientific) were added to 80,000 cells per well in a 6-well dish. These volumes were altered proportionally (according to the above ratio) to accommodate different scales of transfection. To perform siRNA transfections, 5 μl of RNAiMAX (ThermoFisher Scientific), 500 μl of OptiMEM and 5 ul of 20 μM siRNA stock were added to a sub-confluent dish of cells (80,000 cells per well in a 6-well dish). Cells were incubated for 48 hours post-transfection prior to experiments. Control siRNA (5′-CGUUAAUCGCGUAUAAU ACGCGUAT-3′), LIP1 siRNA (5’-ACAAUGCACUGACCGCCUUAGACAG-3’) and TMEM192 siRNA (5’-ACUAUUUCAAGCCUAGAAGAAAUTG-3’) were purchased from Integrated DNA Technologies (IDT, Coralville, IA). Stable expression of STK11IP-GFP was achieved by clonal isolation of cells that had been transfected with the pEGFP-N1-STK11IP plasmid and selected with 500µg/ml G418 (ThermoFisher Scientific).

### Plasmids

For the STK11IP-GFP fusion the desired sequence was amplified by PCR from mouse brain cDNA, digested with HindIII and KpnI and ligated into the pEGFP-N1 plasmid. Human TMEM192 cDNA was likewise amplified by PCR, digested and cloned into the HindIII and KpnI sites in pEGFP-N1. Plasmids for HA-tagged versions of these proteins were generated by site directed mutagenesis which replaced the GFP sequences with HA. TMEM192 deletion mutants were PCR amplified from original plasmid and ligated into HindIII and KpnI-digested pEGFP-N1 or pEGFP-C1 plasmids. STK11IP deletion mutants were PCR amplified from original plasmid and cloned into pcDNA5 FRT/TO (linearized by PCR) by Gibson Assembly. Oligonucleotide primers and cDNA sequences used to generate these plasmids are listed in Table S1.

### Immunoblotting and Co-Immunoprecipitation

Cells were lysed in Tris-Buffered Saline (TBS) + 1% Triton X-100 with EDTA and protease and phosphatase inhibitor cocktails (Roche Diagnostics, Florham Park, NJ). To remove insoluble materials, lysates were centrifuged for 6 minutes at 14,000 rpm. Protein concentrations were measured via Bradford assay prior to denaturation with Laemmli buffer and incubation for 5-minutes at either 55C or 95C. Immunoblotting was performed with 4-15% gradient Mini-PROTEAN TGX precast polyacrylamide gels and nitrocellulose membranes (Bio-Rad, Hercules, CA). Membranes were blocked with 5% milk in TBS with 0.1% Tween 20 (TBST) buffer and then incubated with antibodies in 5% milk or bovine serum albumin in TBST. Antibodies used in this study are summarized in the Table S2. Horseradish peroxidase signal detection was performed using chemiluminescent detection reagents (ThermoScientific) and the Versadoc imaging station (Bio-Rad). FIJI/ImageJ was used to analyze the results and measure band intensities (Schindelin et al., 2012).

For immunoprecipitations, cleared lysates containing GFP-tagged proteins were added to anti-GFP agarose beads (GFP-Trap, ChromoTek, Germany) and incubated while rotating end over end for an hour at 4°C. Beads were the washed 5 times with cold lysis buffer and samples were eluted by addition of a 5x buffer containing 1.2 M β-Mercaptoethanol and 10% sodium dodecyl sulfate (SDS). Where indicated, the immunoblotting techniques described above were then used to analyze these samples.

### Immunofluorescence and Microscopy

For imaging experiments, cells were grown on 12-mm No. 1 glass coverslips (Carolina Biological Supply) and fixed with 4% paraformaldehyde (PFA; Electron Microscopy Sciences, Hatfield, PA) in 0.1M sodium phosphate buffer (pH 7.2) for 30 minutes by dropwise addition of 1 volume of 8% paraformaldehyde in 0.1M phosphate buffer to cells on coverslips in the growth medium. After rinsing with PBS, cells were permeabilized with PBS + 0.2% Triton X-100 for 10 minutes and then blocked for 30 minutes in blocking buffer (5% normal donkey serum/PBS/ 0.2% Triton X-100). Primary antibody incubations were carried out overnight at 4°C in blocking buffer. After washing 3x with PBS+0.2% Triton X-100, samples were incubated in secondary antibody in blocking buffer for thirty minutes at room temperature. Cells on coverslips were then washed 3x with PBS+0.2% Triton X-100 before mounting on slides with Prolong Gold Mounting media (Invitrogen, Carlsbad, CA). Antibodies used in this study are summarized in the Table S4.

For live cell imaging, cells were grown on MatTek dishes (MatTek Corporation, Ashland, MA) and analyzed via spinning disc confocal microscopy at room temperature in a buffer that contained: 136 mM NaCl, 2.5 mM KCl, 2 mM CaCl2, 1.3 mM MgCl2 and 10 mM Hepes, 0.2% Glucose, 0.2% BSA pH 7.4 (Brown et al., 2000). Our spinning disc confocal microscope consisted of the UltraVIEW VOX system (PerkinElmer, Waltham, MA) including the Ti-R Eclipse Nikon inverted microscope (equipped with a 60x CFI Plan Apo VC, NA 1.4, oil immersion) and a CSU-X1 (Yokogawa) spinning disk confocal scan head and Volocity software. Images were acquired without binning with a 14-bit (1,000 × 1,000) EMCCD (Hamamatsu Photonics) and subsequently processed with ImageJ.

### Mass Spectrometry

Two 150-mm dishes of control HeLaM cell and HeLaM cells stably-expressing STK11IP-GFP were lysed, cleared and immunoprecipitated by incubation with anti-GFP beads (ChromoTek). After the immunoprecipitation, the samples were washed twice with TBST and four times with PBS. Samples were eluted in 8M urea and 25 mM Tris. Proteins were digested with endoproteinase LysC and subjected to reverse phase chromatography, mass spectrometry and data analysis as described previously (Frohlich et al., 2013; Hubner et al., 2010).

### Statistical Analysis

Data were analyzed using Prism (Graphpad software) and tests are denoted in figure legends. All error bars represent standard deviations. Data distribution was assumed to be normal, but this was not formally tested.

## Acknowledgements

This research was supported by NIH grants GM105718 and AG062210 to SMF. BA was supported by a National Science Foundation Graduate Research Fellowship (DGE1752134). Agnes Roczniak-Ferguson, Matthew Schrag, Pamela Torola, Tobi Walther and Constance Petit contributed to early phases of this project. The authors do not have any conflicts to declare.

## Author contributions

BA and SMF designed experiments. BA performed all experiments. FF contributed specifically to the mass spectrometry experiment. BA and SMF prepared the manuscript.

## TABLES

**Table S1:**
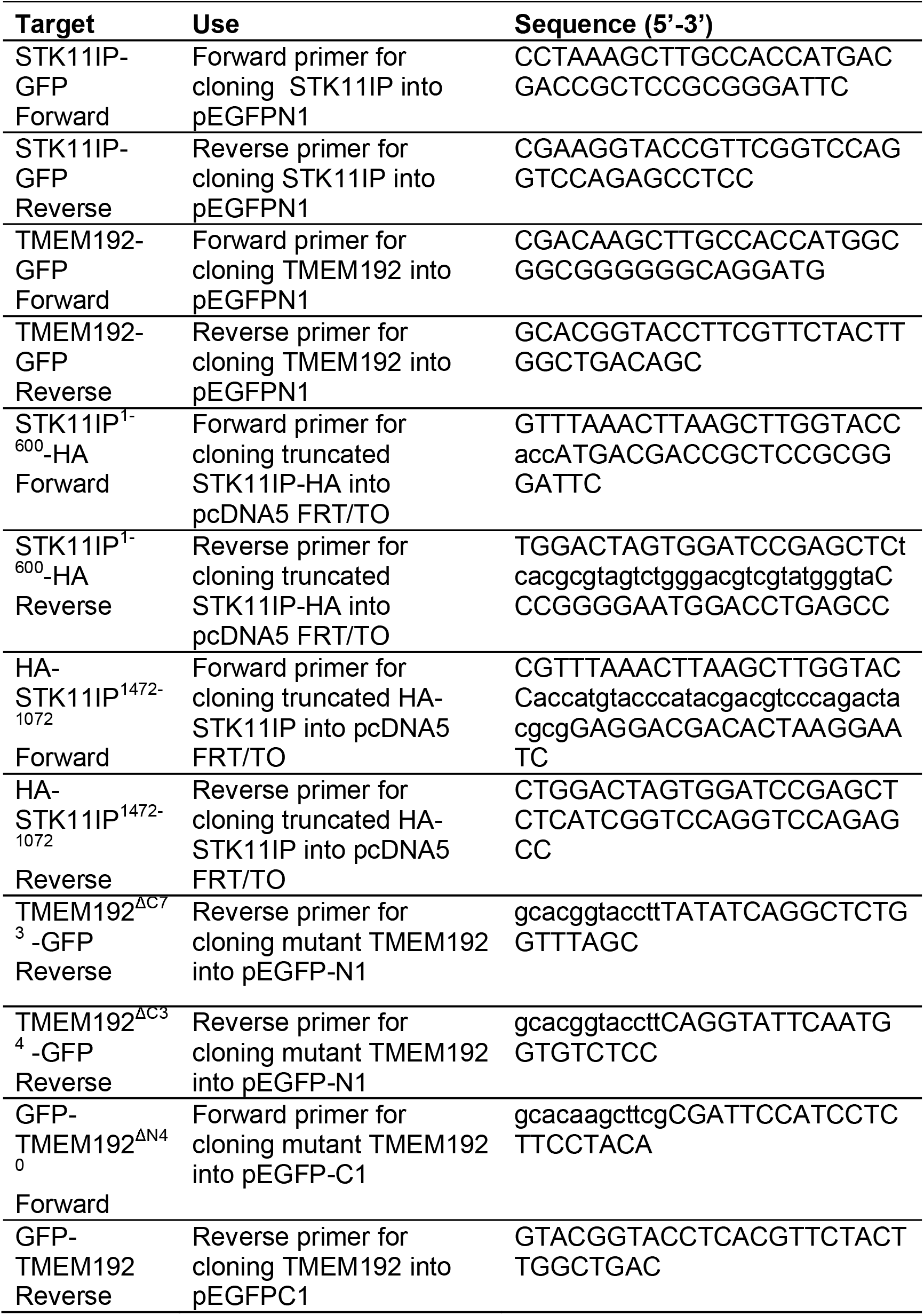
Summary of Oligonucleotides used in Plasmid Construction.

**Table S2:**
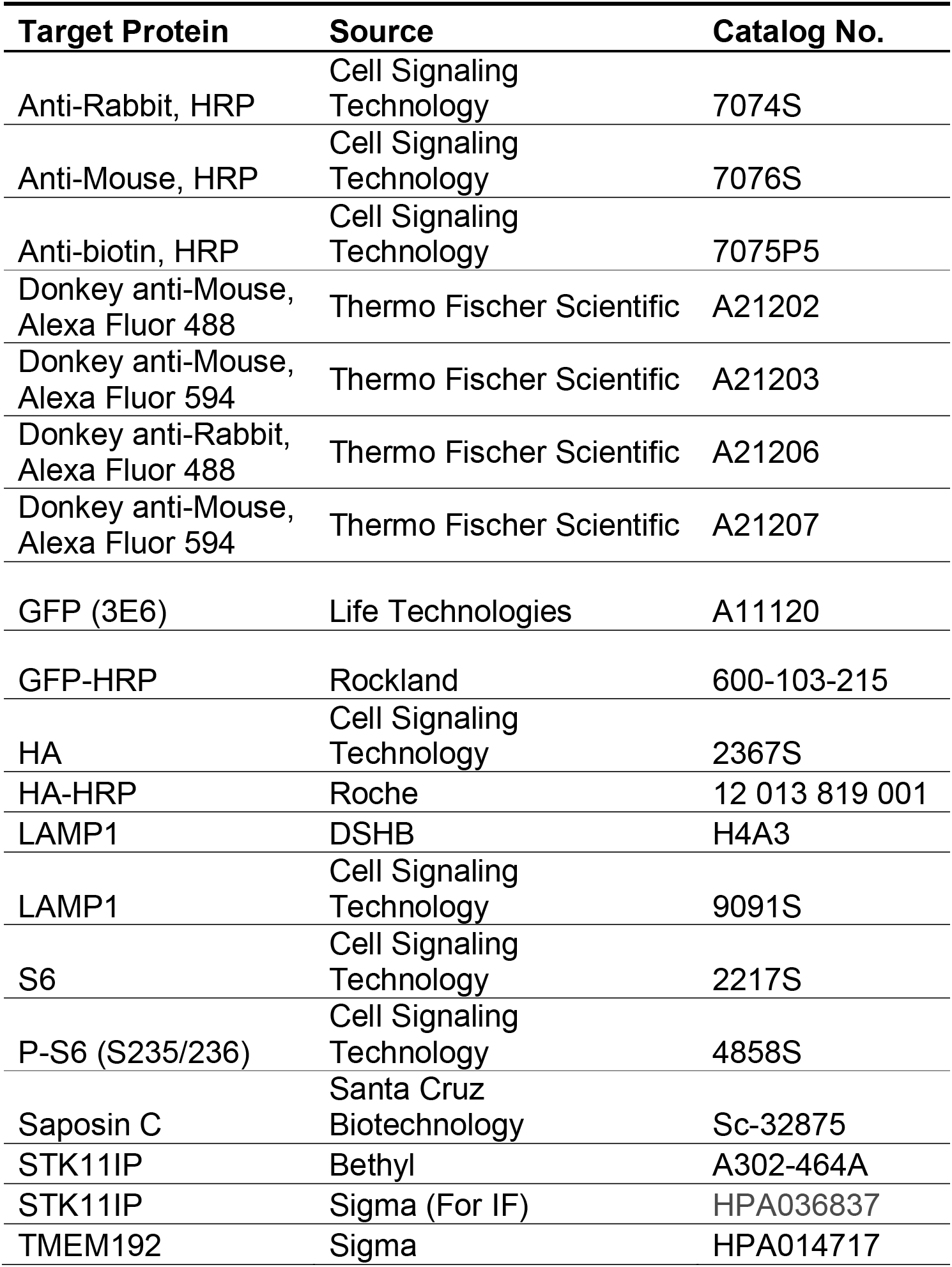
Summary of Antibodies.

